# Implicit vs. explicit choice promotes the combination vs. selection of sensorimotor memories under contextual uncertainty

**DOI:** 10.64898/2026.05.26.727900

**Authors:** Anvesh S. Naik, Sabyasachi Shivkumar, Carlos A. Velázquez-Vargas, James N. Ingram, Máté Lengyel, Daniel M. Wolpert

## Abstract

Skilled action requires expressing motor memories as appropriate for the current context, but context is often uncertain. Theoretical models make conflicting proposals about memory expression under contextual uncertainty, predicting belief-weighted combination of memories versus the selection of the most probable memory. We tested these predictions by training human participants to reach in two opposing force fields cued by the direction of a random dot motion stimulus whose coherence varied. When participants moved before reporting dot direction, adaptation scaled with coherence: low-reliability cues produced partial expression of both memories. Fitting Bayesian observer models to behavior favored belief-weighted memory combination. In contrast, when participants reported their choice before moving, adaptation was independent of coherence and model fits favored categorical memory selection. Thus, sensorimotor memories are expressed as either a probabilistic combination or categorical selection, depending on whether participants’ contextual inference remains implicit or is made explicit at the time of memory expression.

## Introduction

Skilled behavior requires more than learning a single mapping from sensory states to motor commands. The same action can demand different dynamics depending on the object, goal or environment, and the nervous system must therefore store multiple motor memories and express them as appropriate to the current situation. A central proposal is that memories are organized by latent contexts that determine which memories should be updated and recalled [1–9]. Sensory cues, such as an object’s appearance or location, provide evidence for these contexts and can reduce interference between otherwise competing motor memories [10–13].

In everyday behavior, however, contexts are rarely certain. Ambiguous sensory evidence creates a problem for memory expression: should the motor system hedge by partially expressing several plausible memories, or should it commit to one memory and act as if the context had been resolved? One line of work suggests the former [2, 4, 6, 7, 14–16]. In this view, the brain maintains beliefs over possible contexts, and motor output reflects these beliefs. Under uncertainty, the optimal strategy is to form a posterior-weighted combination of the memories associated with each context. We refer to this policy as Memory Combination (*MC*): when evidence is strong, one memory dominates; when evidence is weak, competing memories are blended. This account predicts that memory expression should vary smoothly with cue reliability.

A different prediction follows from work on perceptual decision-making and conditioned perception. Sensory beliefs are hypothesized to be represented probabilistically during deliberation, but once an observer makes a categorical choice, subsequent estimates and actions are be conditioned on that choice [17, 18]. Such commitment promotes internal consistency by treating the chosen hypothesis as the relevant one. Applied to memory expression, this implies Memory Selection (*MS*): after a context has been explicitly chosen, the system should express the memory associated with that choice, with little further dependence on the reliability of the sensory cue. Indeed, a line of previous theories proposed such an all-or-none mechanism for memory selection [5, 8, 9].

We tested whether contextual uncertainty leads to memory combination or memory selection in the case of motor memories, and how this depends on whether, at the time of action, an explicit decision is made about the current context. Participants reached through one of two opposing velocity-dependent force fields. On each trial, the direction of a random dot motion stimulus specified the force-field context, and dot coherence parametrically manipulated contextual uncertainty. In movement-first experiments, participants viewed the stimulus, moved through the force field, and only then reported the perceived dot direction. In choice-first experiments, participants reported the dot direction before moving through the force field. *MC* predicts coherence-dependent adaptation whereas *MS* predicts choice-consistent adaptation that is insensitive to coherence. By combining model-free analyses with Bayesian observer models, we show that the timing of an explicit choice gates motor memory expression between probabilistic combination (movement-first experiments) and categorical selection (choice-first experiments).

## Results

Participants grasped the handle of a robotic manipulandum, while a monitor–mirror system was employed to project visual stimuli into the plane of hand motion (Fig. 1a). On each trial, participants observed a brief (0.6 s) random dot kinematogram (RDK), the global motion direction of which (left vs. right) specified the direction (clockwise vs. counter-clockwise) of the impending velocity-dependent force field through which they were required to move (i.e. the contextual condition). Dot-motion coherence was parametrically manipulated across trials to systematically modulate the level of contextual uncertainty.

**Fig 1.**
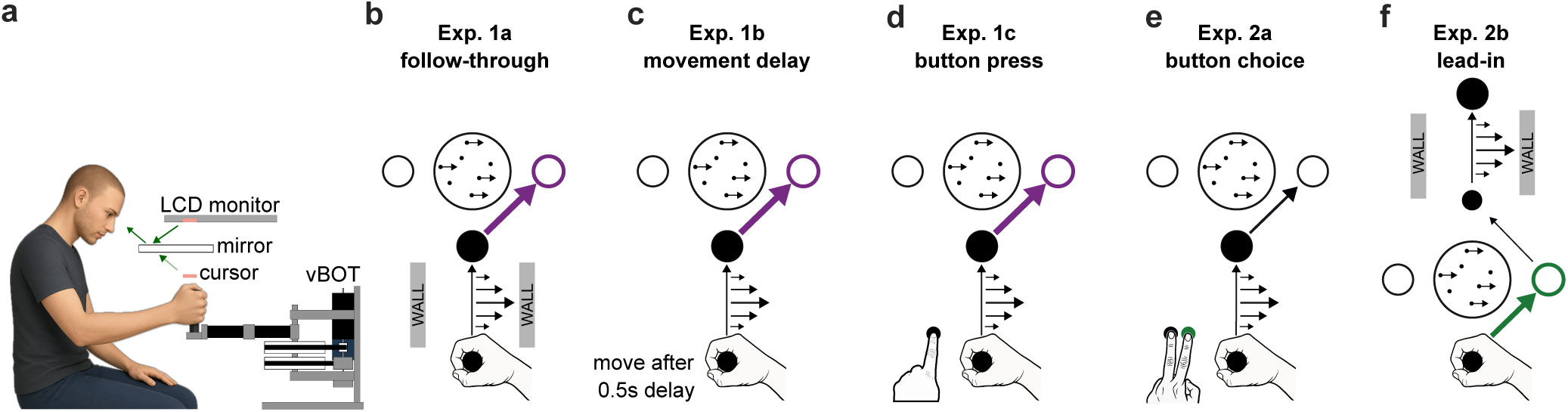
Experimental setup and task paradigms. **a.** Participants grasped the handle of a vBOT robotic manipulandum [19]. A mirror positioned below a downward-facing monitor projected visual stimuli into the plane of the participants’ arm movements. **b.** Follow-through paradigm (Experiment 1a, schematic not to scale). The participant’s hand was initially located at a home position. A random dot kinematogram, with coherent motion either to the left or to the right, was presented for 0.6 s and then extinguished. Participants were subsequently required to reach to an intermediate via-point (large black circle) while experiencing a velocity-dependent force field (horizontal arrows), and then to execute a follow-through movement to one of two possible targets (left or right empty circle). Participants indicated their perceptual judgment of the direction of dot motion (left vs. right) by choosing the corresponding follow-through target. The direction of the force field (clockwise, CW, or counterclockwise, CCW) was deterministically coupled to the direction of motion of the random dot stimulus. **c.** Movement delay paradigm (Experiment 1b), in which participants were instructed to withhold movement initiation for 0.5 s following stimulus offset. **d.** Button-press paradigm (Experiment 1c), in which participants were required to press a single button with their left hand after stimulus offset and prior to initiating the reaching movement with their right hand; the button press was task-irrelevant with respect to target selection. **e.** Button choice paradigm (Experiment 2a), in which participants were required to provide an explicit report of the perceived direction of dot motion by pressing one of two buttons with their left hand after stimulus offset and before initiating the right-hand reaching movement. **f.** Lead-in paradigm in which participants explicitly report the dot motion direction by choosing a left or right via point (empty circles) based on the perceived direction of motion before performing a lead-in movement to a 2nd via point (small black circle) before the movement through the force field. For clarity of illustration, passage walls, which were present in all experiments, are omitted from schematics **c**–**e**. Purple and green indicates movement-first and choice-first decisions, respectively in which in which an explicit decision report was made after (purple) or before (green) the movement through the force field.

### Implicit vs. explicit choices lead to coherence-dependent vs. -independent motor adaptation

We conducted two primary experiments to investigate how contextual uncertainty influences the expression of motor memories. The experiments differed in the timing of participants’ explicit reports of their decision of stimulus direction – either prior to or following movement execution in the force field. As participants were trained with stimulus direction being deterministically coupled to the sensorimotor context of their movement, a decision about stimulus direction implied a decision about sensorimotor context.

In the first experiment (Exp. 1a, follow-through), participants first viewed the visual motion stimulus, then executed a reaching movement through the force field, and only afterward reported their perceived direction of dot motion by moving to a corresponding follow-through target (Fig. 1b). In the second experiment (Exp. 2a, button choice), a separate cohort of participants reported the perceived motion direction using a button box operated with the left hand before initiating the movement through the force field and the subsequent follow-through movement (Fig. 1e). We assessed the influence of contextual uncertainty on the expression of motor memory by varying the strength of the RDK motion and measuring adaptation on occasional channel trials.

After the familiarization phase (hand paths shown in Suppl. Fig. 1), participants were initially trained on the two force fields using exclusively the highest-coherence motion stimuli (*C* = 1, sign indicates motion direction), a condition in which contextual uncertainty is minimal. Over the course of training, movement paths became progressively straighter in both experiments (Suppl. Fig. 1). For a force field-specific measure of adaptation, not confounded by generic changes in stiffness, and to avoid potential bias in follow-through choice that could arise from experiencing the force field, we restricted all subsequent analyses of choices and motor adaptation to occasional channel trials [20, 21] (see Methods). Adaptation during training diverged systematically as a function of motion direction, as expected for two opposing force fields coupled to different follow-through targets [11, 12] (Fig. 2a, columns 1 and 4; adaptation of last five trials: *t*_9_ = 6.5, *p* = 1.1 10^−4^ in Exp. 1a; *t*_9_ = 10.6, *p* = 2.2 10^−6^ in Exp. 2a). These results indicate that, under unambiguous contextual evidence, participants acquired and expressed distinct motor memories for the two opposing force fields in both experiments.

**Fig 2.**
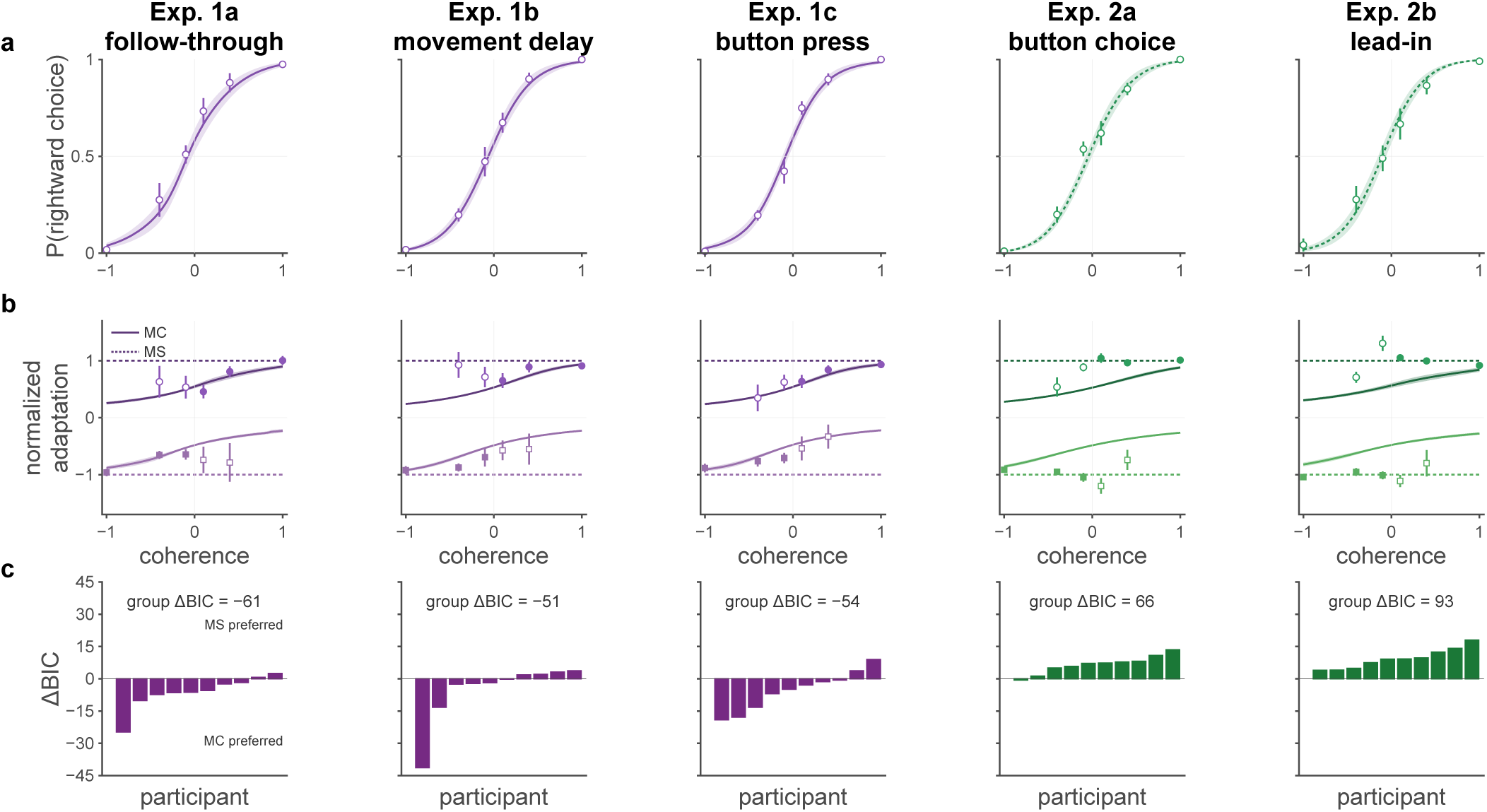
Implicit vs. explicit choices lead to coherence-dependent vs. -independent motor memory expression. Results for each experiment (columns). **a.** adaptation measured on channel trials in familiarization and training phases. Data shows mean *±* s.e.m. for channel trials split by dot motion direction (positive going traces for rightward motion). **b** Probability of a rightward choice on channel trials. Solid lines show the across-subject mean probit fit and shaded regions indicate s.e.m. **c** mean *±* s.e.m. probit slopes with individual slopes (points) for the conditions. All pairwise comparisons were performed and none were significant. **d**. normalized adaptation slope comparison (mean *±* s.e.m and individual participants) between the five experiments. The data is split by choice (dark for rightward). Both the data and regression fits have been normalized (see Methods). Both correct (filled) and incorrect (unfilled) choice trials were included. Data points shown are mean *±* s.e.m. (some error bars are smaller than the points). For clarity, data for incorrect choice trials for *C* = *±*1 are not shown as very few participants had such errors, but all trials were used during model fitting). **e.** mean *±* s.e.m. adaptation slopes with individual slopes (points) for the conditions. All pairwise comparisons were performed (\**p <* 0.05, \*\**p <* 0.01 and \*\*\**p <* 0.001, *p >* 0.05 not shown; two sample t-tests). Note here and in other figures the data and models are color coded by whether an explicit decision report was made after (purple) or before (green) the movement through the force field.

During the testing phase, we also included trials with lower coherence. Psychometric curves (Fig. 2b) showed, as expected [22] that the probability of rightward choice increased with coherence. However, there was no significant difference in choice sensitivity (slopes of individual probit fits) between the two experiments (two-sample *t*-test: *t*_9.8_ = 1.08, *p* = 0.31).

Previous work on the same perceptual decision making task showed that participants’ uncertainty is commensurate with their psychometric performance [23, 24]. Given the deterministic coupling of stimulus direction and force-field direction in our experiment, higher uncertainty about the stimulus on low coherence trials also implies higher uncertainty about the upcoming sensorimotor context (i.e. the direction of the force field). Thus, we examined how the expression of motor memories in adaptation varied with coherence.

We considered two hypotheses as to the effect of contextual uncertainty on the expression of sensorimotor memories when different memories were created for different contexts. The normatively appropriate way to take contextual uncertainty into account is to perform memory combination: expressing a combination of memories, each weighted by the probability with which the current context is believed to be the context for which it was created. The alternative is memory selection: only expressing the memory which was created for the currently most probable context. At the highest coherence levels, both these hypotheses predict that a single memory (that created for the force field that has been coupled to the unambiguously perceived stimulus) is expressed, resulting in high adaptation. However, at lower coherence levels, their predictions diverge. Memory selection predicts that, once conditioned on the perceptual decision (reflecting the most probable context), memory expression should be independent of coherence. In contrast, memory combination predicts that, even when conditioning on the perceptual decision, memory expression should vary with coherence because the underlying context probabilities that are used to weight the memories are not binary and depend parametrically on coherence. Specifically, when coherence decreases, the incorrect memory is assigned an increasing probability, making adaptation smaller.

To test whether memory combination or selection best accounted for movements, we first performed a model-free analysis, in which we regressed adaptation against coherence. In the follow-through experiment, adaptation was significantly modulated by coherence (Fig. 2d, column 1; *β* = 0.17 ± 0.03, *t*_9_ = 5.1, *p* = 6.8 × 10^−4^, BF = 56.2). In contrast, in the button choice experiment, this modulation remained non-significant (Fig. 2d, column 4; *β* = 0.02 ± 0.02, *t*_9_ = 1.0, *p* = 0.32, BF = 0.48), and there was a significant difference between the two experiments (Fig. 2e; two-sample *t*-test: *t*_14.5_ = 3.8, *p* = 0.0017). These results provide evidence for memory combination in the follow-through experiment, in which the choice of context remained implicit during movement through the force field, and memory selection in the button choice experiment, in which context choice was explicitly reported before movement.

### Bayesian observer models

Our model-free analyses suggested it was possible to distinguish between memory combination and selection. However, they involved an arbitrary assumption that adaptation would be linearly related to coherence (with a positive or a zero slope for the two hypotheses), while the data suggested there may be a non-linear relationship (Figure 2b). Thus, to adjudicate between memory combination and memory selection in a more principled way, we developed Bayesian observer models for them (Fig. 3, Methods). These models shared a common part that performed Bayesian inference over context as a binary latent variable by combining prior expectations about stimulus direction with stimulus evidence (Fig. 3 top box), of which the noise varied parametrically with coherence. The resulting posterior distribution (Fig. 3 middle box) was used to drive both the perceptual decision and the movement of the observer (Fig. 3 bottom box). Perceptual decisions (expressed by the choice of end target in follow-through, or by pressing the appropriate button in button choice) always reflected the stimulus direction that was assigned the highest probability under the observer’s posterior distribution. The observer also had two motor memories, one for each force field context, and the two models differed in how they used the posterior over contexts to drive movement based on these memories. The memory combination model expressed a combination of the two memories, each weighted by the probability of its corresponding context under the posterior distribution. The memory selection model expressed a single memory, the one that had maximal probability under the posterior (i.e. the one that corresponded to the perceptual decision). Both models allowed additional noise in the adaptation they expressed. We fit both models to each participant’s data for the channel trials of the testing phase, simultaneously fitting both perceptual decisions and motor adaptation (Fig. 4a,b, see Methods). Both models had the same number (5) of parameters that included a bias parameter for the strength of prior expectations of a leftward v.s. rightward stimulus, the noise variance on sensory observations of the stimulus, the level of adaptation stored in each of the two motor memories, and the noise variance on motor adaptation output.

**Fig 3.**
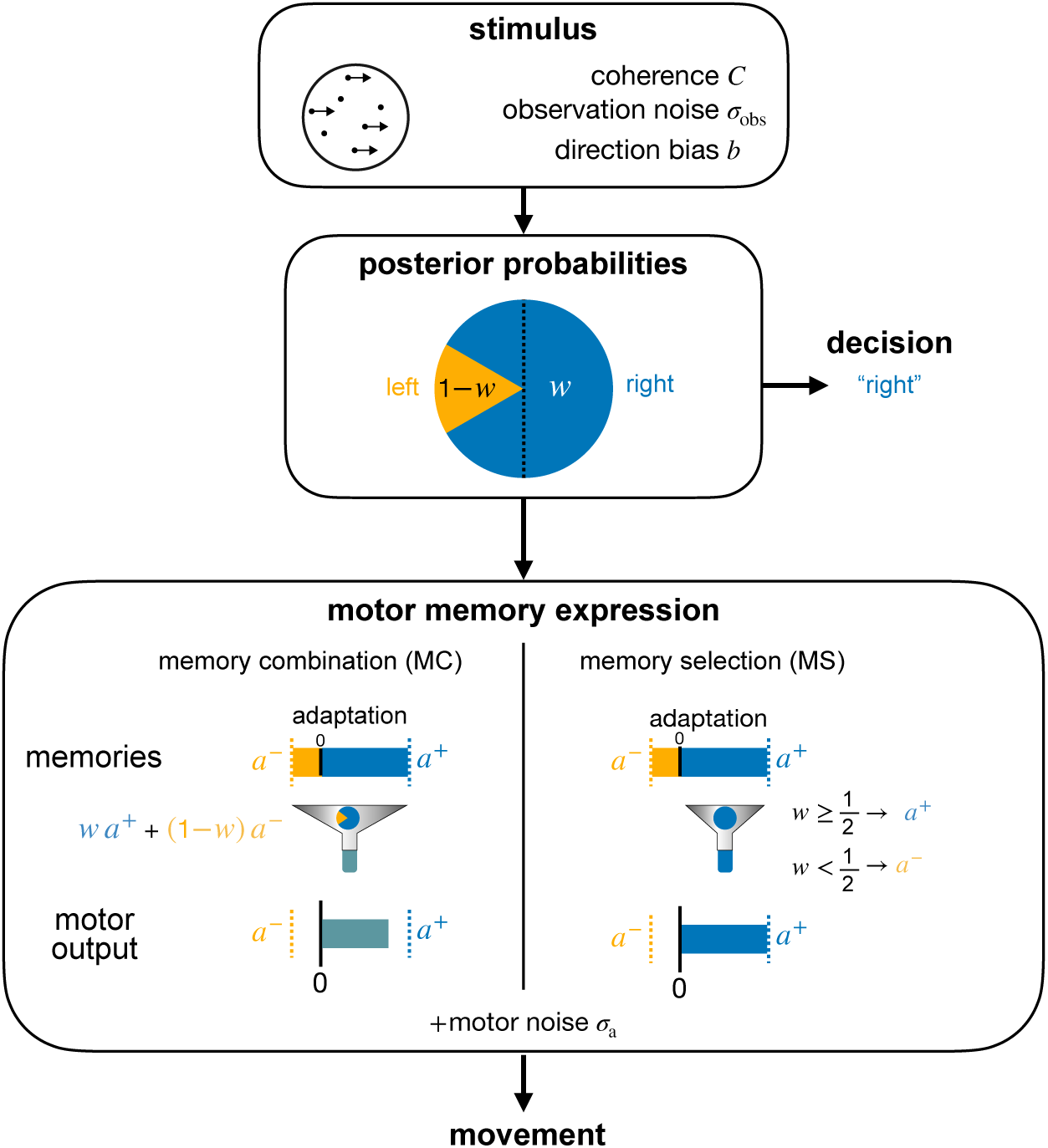
Bayesian observer model. On each trial the stimulus (RDK; top box) has a coherence *C* (varied across trials) and the participant views the stimulus with observation noise (*σ*_obs_) and, possibly, a motion direction bias (*b*). On each trial the participant makes an inference (middle box) reflected in the posterior probabilities (pie chart) over motion directions (*w* for right and 1 *−w* for left). The decision is based on the highest posterior (in this example right). The motor memory expression (bottom box) that leads to movement depends on whether the observer uses memory combination (*MC*) or selection (*MS*). Under memory combination the memories (*a*^+^ and *a*^−^; shown here with asymmetric learning) are combined based on the posterior (pie chart) to produce a weighted output. Under memory selection only the memory with the highest posterior is expressed. In both cases the expressed adaptation is also corrupted by motor noise (*σ*_a_).

**Fig 4.**
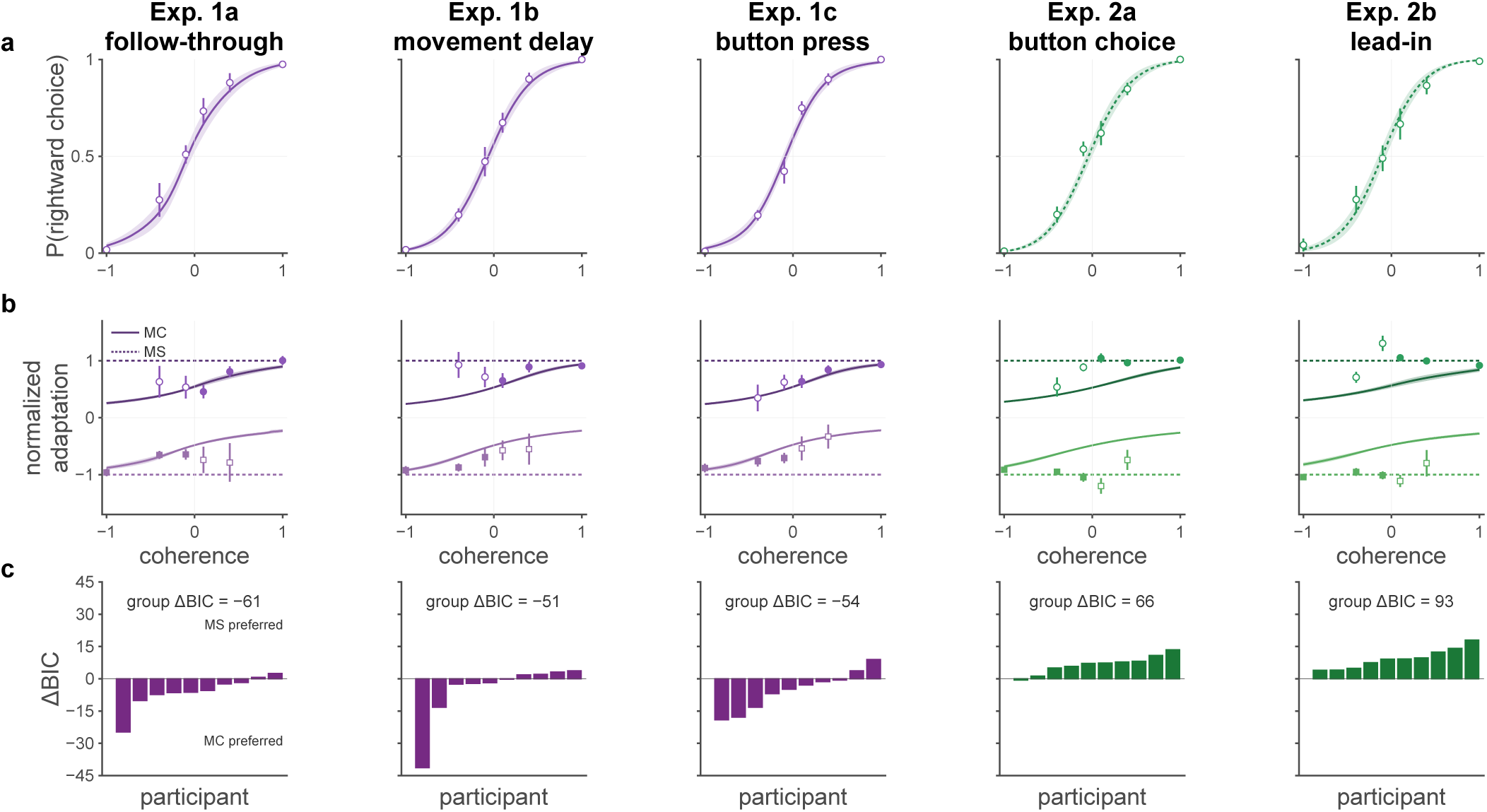
Implicit choices lead to motor memory combination but explicit choices leads to memory selection under contextual uncertainty. Bayesian observer model fit to all experiments (columns). **a–b** probability of a rightward choice and adaptation on channel trials. Data points and model (solid and dotted lines) shown are mean *±* s.e.m.(some error bars are smaller than the points). For **b** All correct (filled) and incorrect (unfilled) choice trials were used for fitting but for clarity, data for incorrect choice trials for *C* = *±*1 are not shown as very few participants had such errors. Memory combination (*MC*) and memory selection (*MS*) fits are shown with solid and dotted lines, respectively. For choice fits (**a**), only the best model fits are shown (*MC* for first 3 columns and and *MS* for last two columns) **c.** ΔBIC for model comparison for individuals participants with negative in favor of *MC*.

Model fits for the two experiments are shown in Fig. 4a,b (columns 1 and 4) and individual participant fits are shown in Suppl. Fig. 2. For the follow-through experiment, model comparison provided strong support for memory combination both at the group level (ΔBIC = −61) and at the individual participant level (Fig. 4c, column 1; ΔBIC *<* 0 for 8/10). Conversely, for button choice, model comparison provided strong support for memory selection both at the group level (ΔBIC = 66) and at the individual participant level (Fig. 4c, column 4; ΔBIC *>* 0 for 9/10). These model-based analyses are therefore consistent with the model-free regression results reported above. In follow-through – where choices were reported at the end of the movement – memory combination provided a superior account of trial-by-trial adaptation, capturing the graded decrease in adaptation as contextual uncertainty increased. In contrast, in button choice, where choices were explicitly made before movement, memory selection outperformed memory combination, producing the flat adaptation profile across coherence.

These combined behavioral and computational findings demonstrate that under contextual uncertainty participants combine the memories for the force fields, weighting them by their belief that they are appropriate for the current situation (Exp. 1a). However, when an explicit decision is reported (Exp. 2a), they no longer use uncertainty to blend the memories but instead select the memory consistent with their report. Therefore, the order of explicit decision reporting may gate whether learned motor memories blend or whether only one is chosen and fully expressed.

### Time delays and motor actions in the absence of explicit choice lead to memory combination

The main experiments demonstrated that contextual uncertainty influences motor-memory expression in distinct ways depending on when an explicit choice is reported relative to memory expression. However, these differences could, in principle, arise from other aspects of the task design rather than the temporal order of explicit choice. In the button choice experiment, there was an additional delay between stimulus offset and movement onset due to the button press. This delay might allow participants to internally commit to a target before initiating the movement through the field. Moreover, this experiment required an additional motor action (the button press) prior to movement, which could itself influence subsequent memory expression.

To address these potential confounds (and replicate the key results), we conducted two variants of the follow-through paradigm. These experiments isolated the effects of temporal delay and additional motor action while preserving the overall task structure. In the movement delay paradigm (Exp. 1b), participants were required to wait for 0.5 s after stimulus offset before initiating the movement through the field (Fig. 1c). This manipulation isolated the effect of the temporal delay present in the button choice paradigm while preserving the follow-through structure. In the button press paradigm (Exp. 1c), participants were required to press a single button immediately after stimulus offset, prior to initiating the movement through the field (Fig. 1d). Crucially, this button press was irrelevant to target choice and served only to replicate the additional motor action required in the button choice paradigm, without requiring an explicit decision before movement.

Suppl. Fig. 1 illustrates the progression of paths in these experiments. Again, movements became progressively straighter over the course of training, indicating adaptation to the opposing force fields. In both experiments, adaptation measured on channel trials (Fig. 2a columns 2 and 3) differed significantly for the two coherences (Exp. 1b: *t*_9_ = − 5.9, *p* = 2.3×10^−4^; Exp. 1c: *t*_9_ = − 7.4, *p* = 4.0×10^−5^). Pairwise comparisons of the slopes of the psychometric functions revealed no significant differences across experiments (Fig. 2c).

The adaptation showed coherence-dependence, as revealed by a positive slope in the model-free regression analysis (Fig. 2d–e columns 2 and 3; Exp. 1b: *β* = 0.12 ± 0.02, *t*_9_ = 5.3, *p* = 4.8 × 10^−4^, BF = 76.0; Exp. 1c: *β* = 0.17 ± 0.04, *t*_9_ = 3.9, *p* = 0.0034, BF = 14.9). These slopes did not significantly differ from one another (Fig. 2e; Exp. 1b vs. 1c: *t*_13.7_ = −1.0, *p* = 0.34) or from the follow-through experiment (Exp. 1b vs. 1a: *t*_15.9_ = −1.2, *p* = 0.27; Exp. 1c vs. 1a: *t*_16.9_ = 0.04, *p* = 0.97), while significantly differing from the button choice experiment (Exp. 1b vs. 2a: *t*_17.5_ = −3.4, *p* = 0.003; Exp. 1c vs. 2a: *t*_12.5_ = 3.2, *p* = 0.007).

Model-based analyses were also consistent with these results (Fig. 4 columns 2 and 3; individual fits in Suppl. Fig. 2), providing strong support for the memory-combination model at the group level (Exp. 1b: ΔBIC = −51; Exp. 1c: ΔBIC = −54). At the level of individual participants, however, there was a weaker support for Exp. 1b (ΔBIC *<* 0 for 6/10), whereas a strong support for Exp. 1c (ΔBIC *<* 0 for 8/10). Thus, despite controlling for temporal delay and a motor action prior to movement through the field, these two control experiments suggest that contextual uncertainty leads to graded memory expression when an explicit choice is only reported after memory expression.

### Reporting explicit choice by lead-in movement before action leads to memory selection

Pressing buttons for reporting choice in the button choice paradigm required qualitatively different movements, and with a different hand, than those controlling the cursor in the reaching task. We wondered whether these differences were necessary for the explicit reporting of choice to result in the selection rather than combination of memories. Thus, we conducted a final lead-in experiment (Exp. 2b; Fig. 1f). In this experiment, participants explicitly committed to a choice before the movement through the force field by initiating a reach directly toward their chosen lateral target after viewing the stimulus. They then executed a lead-in movement to a via-point, followed by the movement through the field. This design preserved the central conceptual feature of button choice – an explicit decision made before the field movement – while using the same reaching paradigm with the same hand as in the follow through paradigm.

As in the main experiments, paths became progressively straighter (Suppl. Fig. 1) and adaptation curves clearly diverged (Fig. 2a, column 5; *t*_9_ = 12.1, *p* = 7.2 × 10^−7^). Again, the slope of the psychometric curve did not differ significantly from any of the other experiments (Fig. 2c).

The model-free regression analysis (Fig. 2d-e, column 5) revealed slopes that did not significantly differ from zero (*β* = 0.03 0.02, *t*_9_ = − 1.6, *p* = 0.15, BF = 0.78) or from those found in the button choice experiment (Exp. 2b vs. 2a: *t*_17.8_ = − 1.8, *p* = 0.086), while significantly differing from the slopes found in movement-first experiments (Exp. 1a vs. 2b: *t*_13.5_ = 5.2, *p* = 0.0001; Exp. 1b vs. 2b: *t*_16.8_ = 5.2, *p* = 0.0001; Exp. 1c vs. 2b: *t*_11.8_ = 4.2, *p* = 0.0012).

The model-based analysis (Fig. 4c column 5) also provided strong support for memory selection both at the group level (ΔBIC = 93) and at the level of individual participants (ΔBIC *>* 0 for 10/10).

## Discussion

We show that the same contextual uncertainty can lead to two different forms of motor memory expression. When participants moved before reporting the direction of the contextual cue, adaptation declined smoothly as cue reliability decreased, and Bayesian observer models favored belief-weighted memory combination. When participants reported their choice before moving, either with a button press or with a lead-in reach, adaptation was largely independent of cue reliability, and models favored categorical memory selection. Control experiments showed that this dissociation was not explained by the additional delay before movement or by the presence of an extra pre-movement action. Rather, the critical factor was whether participants had explicitly reported a context before the motor memory had to be expressed.

One potential confound in our analyses could be caused by decision noise. Both our model-free and model-based analyses assumed that participants’ choice deterministically reflects the stimulus direction that is assigned higher posterior probability at the time of choice. This is a standard assumption in the bulk of work on perceptual decision making with the RDK stimuli [22]. Nevertheless, it is possible that participants have finite and graded amounts of decision noise, such that the more even the posterior probabilities corresponding to the two stimulus directions are (i.e. the higher the level of contextual uncertainty is), the more probabilistic choices become – in line with often described forms of probability matching behavior in other paradigms [25]. In this case, even if motor memory expression was purely based on the most probable context (i.e. *MS*), conditioning motor adaptation on participants’ choice (as we did) could still mix data from trials in which either context was inferred most probable, and thus make motor adaptation appear as a graded function of contextual uncertainty, reminiscent of *MC*. However, it is unlikely this confound could account for our results. In particular, it could only account for the differences in memory expression we found in Experiments 1a–c versus 2a–b if our experimental manipulations had affected the level of decision noise. The near-identical slopes of the psychometric curves we found across all our experiments (Fig. 2c) seem incompatible with this account.

A previous study also required participants to adapt to force fields depending on an RDK stimulus of which the coherence was manipulated [26]. However, in that study, the force field experienced by participants depended on the coherence (rather than the direction) of the RDK stimulus, which varied categorically between two extreme levels. Thus, there was no uncertainty about the force field on any given trial, and consequently no opportunity to study the effect of contextual uncertainty on memory expression.

Our findings extend contextual inference accounts of motor learning. Multiple internal model theories propose that separate memories are learned for different contexts and selected according to the inferred state of the environment [1–5]. The theoretical framework of contextual inference formalizes this idea by treating context as a latent variable and predicting that, under uncertainty, motor output should reflect posterior beliefs over contexts [4, 6, 7, 15, 16]. Previous experimental evidence was broadly consistent with this idea, but without a direct experimental manipulation of contextual uncertainty it remained circumstantial [7]. By manipulating contextual uncertainty via cue coherence, our movement-first experiments provide direct behavioral evidence for memory combination: when the contextual cue was unreliable, participants did not simply express the memory associated with their eventual categorical report, but instead expressed a graded mixture of the two learned force-field memories.

In contrast, the choice-first experiments show that probabilistic expression is not inevitable. This result speaks directly to a specific point in a previous contextual inference-based theoretical account of explicit and implicit visuomotor learning [7]. When applied to modeling visuomotor rotation paradigms [27, 28], the theory was only able to account for the empirical data by assuming that the form of memory expression depended on the way participants reported their aiming direction. When this was purely implicit in the way they moved their finger, the theory predicted the standard posterior probability-weighted expression of the memories of two opposing perturbations. When an explicit aim judgment was solicited before motor output, the theory predicted that movement would reflect the memory associated with the most probable context (i.e. memory selection), rather the posterior-weighted combination of memories. While these predictions qualitatively matched the data, the experimental paradigms also did not directly control contextual uncertainty, and thus provided only indirect evidence for the key role of contextual uncertainty in these effects. Our experiments provide a direct test of that previously inferred distinction.

More broadly, our results support the view that decision-making and motor control are coupled rather than serial processes [29]. Sensorimotor circuits can represent multiple potential actions during deliberation and resolve this competition as commitment approaches [30–32]. Preparatory activity in motor cortex can also occupy output-null states that do not yet drive movement, while later transitions into output-potent activity allow the selected plan to be expressed [33, 34]. Recent population-level studies of perceptual decision-making extend this view by showing that deliberation and commitment correspond to distinct neural states: during deliberation, neural dynamics remain sensitive to incoming evidence, whereas after commitment, dynamics shift into a post-commitment regime in which subsequent evidence has little influence on the chosen outcome [35–37]. Our behavioral dissociation may reflect an analogous transition at the level of learned dynamics: before commitment, competing context-specific memories can jointly influence the motor command, whereas after commitment, expression may be constrained to the selected context. In the case of force-field adaptation, such a transition would have to act on the cortico-cerebellar circuits that support learned compensation. Consistent with this possibility, cerebellar output has been shown to shape cortical preparatory activity during force-field adaptation, and cerebellar signals can carry sensory, choice-, and evidence-related information [38–42]. Our data do not identify the neural mechanism, but they define the behavioral computation that such mechanisms must explain.

In conclusion, contextual uncertainty does not have a single consequence for motor memory expression. When action proceeds without (or before) reporting an explicit choice, as may be the case in most ecologically relevant settings, memories are combined as normatively appropriate for movement, producing graded adaptation. The explicit reporting of choice seems to collapse uncertainty, resulting in memory selection. This is consistent with recent suggestions that communication (or the intent to communicate) transforms the content of internal representations [43]. Our results, together with earlier work on conditioned perception [18], hint at the possibility that a critical aspect of such a transformation may be the collapse of uncertainty.

## Methods

### Participants

A total of 50 participants (21 males and 29 females) aged 18–60 years old (28.9 ± 6.6 years) were recruited. The participants were right-handed according to the Edinburgh handedness questionnaire and reported that they had normal or corrected to normal vision and no prior diagnosis of movement disorder. They were compensated at a rate of $17 per hour. All experiments were conducted in accordance with the Declaration of Helsinki of 1964, following a protocol approved by the Columbia University Institutional Review Board (AAAR9148). Written informed consent was obtained from all participants prior to their participation.

### Experimental apparatus

Experiments were performed using a vBOT planar robotic manipulandum [19] with an integrated virtual-reality display (Fig. 1a). This custom-built, back-drivable device features a low-mass handle and records position and force data at 1 kHz. Participants grasped the handle with their right hand and moved in the horizontal plane. Visual feedback was provided via a computer monitor projected through a horizontal mirror so that all images appeared in the plane of hand movement. Stimulus presentation and task control were implemented using custom MATLAB software (MATLAB 2023a) with Psychtoolbox (version 3.0.19).

### Paradigm

The basic paradigm involved participants making a movement through a force field whose direction was determined by the motion direction of a random dot kinematogram (RDK) displayed prior to movement initiation.

Participants were assigned to one of five experimental groups (n = 10 per group). Four experiments were variants of a follow-through paradigm [11] in which participants moved through the force field before moving to one of two follow-through end targets to indicate the perceived dot motion direction. The fifth group was a lead-in paradigm [44] in which participants moved to one of two lateral targets (to indicate the perceived dot motion direction) before moving through a force field between a via point and the end target (full details below).

For all experiments, a 0.5 cm-radius hand cursor displayed the hand position in the movement plane. In the follow-through experiments, movements were made from a home position (0.25 cm-radius circle, approximately 30 cm from chest) to a via-point (1 cm-radius circle) 10 cm distal to the home position, and then to one of two lateral end targets (1 cm-radius circles) 5 cm distal to the via-point at 45^◦^ (Fig. 1b). Two rectangular visually represented “walls” flanked the path between the home position and the via-point, creating a narrow (6 cm-wide) passage. If the walls were contacted by the hand cursor this led to a miss trial.

In the lead-in experiment (Fig. 1f), the order of via-point and lateral targets was reversed. Participants first moved from the home position to one of two lateral targets and then to the via-point before moving to an end target (circle of 1 cm radius) 10 cm distal to the via-point via a narrow passage flanked by the two “walls”.

Each trial began with the vBOT passively moving the participant’s hand to the home position where it was held stationary and a random dot kinematogram was displayed. The detailed timeline of the rest of the trial depended on the specific experiment (see below).

### Random Dot Kinematogram (RDK)

A dynamic random dot stimulus appeared between the lateral targets. The stimulus was presented within an aperture subtending 2.5^◦^ of visual angle for a duration of 0.6 s and then disappeared. The motion stimulus is described in detail in previous studies [45]. Briefly, three banks of dots were used with dots in each bank displayed for one frame (16.7 ms, 60 Hz refresh) and then three frames later a subset of these dots was displaced in the direction of motion (either left or right on each trial) and the rest of the dots were repositioned randomly. The dot density was 16.7 dots deg^−2^ s^−1^ and displacements were consistent with a motion speed 7.1 deg/s. On each trial, the difficulty of the motion discrimination task was determined by the coherence of the stimulus, defined as the probability that each dot would be displaced as opposed to randomly replaced. The signed coherence (*C*) on each trial was selected from the set of *C* = 0.1, 0.4, 1.0 , where negative coherence corresponds to the leftward motion (direction *D* = left) and positive coherence to rightward motion (direction *D* = right).

### Force fields

On each trial, the vBOT could either generate no forces (null field) or a velocity-dependent viscous curl field (force-field trials) or a force channel (channel trials) between home position and via-point in follow-through experiments or between via-point and end target in the lead-in experiment. All other movements were made in a null field. The velocity-dependent force field (used in many previous experiments [e.g. 46, 47]) is given by

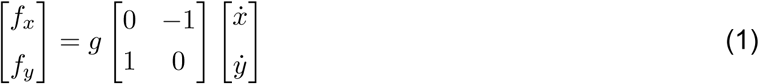

where *f_x_*, *f_y_*, *x* and *y* are the forces and velocities at the robot handle in the *x* and *y* directions (mediolateral and anteroposterior, respectively). The field gain *g* was set to 15 or +15 N s m^−1^, corresponding to a clockwise (CW) and counter-clockwise (CCW) field, respectively. On channel trials, the hand was constrained to move along a straight line between targets. This was achieved by simulating forces associated with a stiff spring and damper, with the forces acting perpendicular to the long axis of the channel. A spring constant of 6, 000 N m^−1^ and a damping coefficient of 5 N s m^−1^ were used. Channel trials allowed the feedforward forces generated by participants to be measured orthogonal to the direction of reach [20, 21]. Crucially, the sign of *g* depended on the direction (sign) of the RDK stimulus (*D* = ±1) and this specified whether the curl field was CW or CCW on a given trial. The association between curl-field direction and stimulus direction was counterbalanced across participants.

### Experimental phases

Each experiment was divided into three phases: familiarization, training, and testing phases.

The *familiarization* phase comprised 54 null-field trials, split into nine blocks with six trials per block. On each trial, the stimulus coherence was drawn randomly without replacement (in a block) from the set of six coherences (see above). Thus, each coherence level appeared once per block. Two randomly selected trials per block were channel trials, and the rest were null-field trials. We paired channel trials with the same magnitude of coherence (*C*) within a block and ensured that both trials did not occur consecutively or as the first or last trial of a block. Across blocks, the channel-trial coherences were balanced so that each coherence level was repeated an equal number of times. At the end of each trial, participants received accuracy feedback: a pleasant chime indicated a correct choice and a harsh tone indicated an incorrect choice.

The *training* phase comprised 324 force-field trials, split into 63 blocks with six trials per block. On each trial, the stimulus coherence was pseudorandomly selected from *C* = ±1 (with equal number across the phase). Within each block, two trials were designated as channel trials, one with *C* = +1 and the other with *C* = −1, and the rest were force-field trials (consistent with the stimulus, see above). As in the familiarization phase, the two channel trials never appeared consecutively or as the first or last trial of a block. At the end of each trial, a neutral tone was presented, irrespective of choice accuracy.

The *testing* phase comprised 216 force-field trials, split into 36 blocks with six trials per block. As in the familiarization phase, the stimulus coherence was drawn randomly without replacement (in a block) from the set of six coherences on each trial. However, instead of using null fields, force fields were paired consistently with the stimulus (see above), with the exception of two trials per block, which were channel trials as in the familiarization and training phases. At the end of each trial, a neutral tone was presented, as in the training phase.

In total, participants performed 594 trials, rest breaks were provided after every 54 trials, and a session lasted 60 to 70 minutes.

We performed five experiments in total that differed in the order of explicit choice reporting relative to movement through the force field. In experiments 1a–c (movement-first), participants first executed the movement through the force field and subsequently reported their perceptual choice by moving to the corresponding end target. In experiments 2a–b (choice-first), participants explicitly reported their choice before initiating the movement through the force field. This manipulation allowed us to test how the order of an explicit decision relative to movement influences motor memory expression.

### Experiment 1a: Follow-through

This experiment employed a variation of a follow-through paradigm (Fig. 1b). Participants were asked to estimate the direction (leftward or rightward) of the dot motion. After the RDK disappeared, an auditory tone cued participants that they could initiate their movement from the home position toward the via-point. Participants were required to initiate movement within 1 s of the cue and, upon reaching the via-point, to continue with a follow-through movement to the lateral end target corresponding to their estimate of the direction of the dot motion. The force field (or channel) was applied between the home position and the via-point. Follow-through movements between the via point and the lateral end targets were always performed in a null field. A trial ended when the final target was reached.

### Experiment 1b: Movement delay

This experiment was identical to Experiment 1a (follow-through), except that there was a 0.5 s delay between the disappearance of the RDK and the auditory cue (Fig. 1c). This delay was introduced in order to control for a critical difference between Experiments 1a & 2a (button choice, see below). Specifically, the button choice in Experiment 2a made the time elapsed between stimulus offset and movement onset (0.65 ± 0.06 s) longer than in Experiment 1a (0.16 ± 0.01 s). Thus, the 0.5 s delay in this experiment was chosen to approximately match the average temporal difference between Experiments 1a & 2a. This allowed us to test whether the differences observed between those experiments were driven not by the order of explicit choice relative to movement, but simply by the additional time available between stimulus offset and movement initiation.

### Experiment 1c: Button press

This experiment was another way to control for the differences between Experiments 1a & 2a. Specifically, it was identical to Experiment 2a (see below), except that participants pressed the same button with their left index finger on every trial, which was thus unrelated to the perceptual decision (Fig. 1d). Compared to Experiment 1b (movement delay), which only controlled for the additional time between stimulus offset and movement initiation, this experiment also controlled for the additional motor action required to press a button. This allowed us to study whether the mere execution of a discrete action prior to movement –rather than the explicit reporting of a perceptual choice – influenced motor memory expression. The time elapsed between stimulus offset and movement onset was 0.45 ± 0.04 s (mean ± s.e.m).

### Experiment 2a: Button choice

This experiment modified Experiment 1a by introducing an explicit choice report before movement (Fig. 1e). After the auditory tone and prior to initiating the reach, participants indicated their estimate of the dot motion direction (and hence which end target they would move to) by pressing the left or right button on a response box with the middle or index finger of their left hand, respectively. The button press was required within 0.5 s of the cue. Following the response, participants executed the reaching movements as described in Experiment 1a. This allowed us to test whether reporting one’s explicit choice before movement influenced the expression of the corresponding motor memory. Although, a participant could in principle report one choice and then execute the opposite follow-through after experiencing the field, in practice such changes of mind were extremely rare, occurring on around 0.1% of trials across participants.

### Experiment 2b: Lead-in

Compared to the experiments using a follow-through paradigm (Experiments 1a–c & 2a), this experiment reversed the order of intermediate and final targets (Fig. 1f). Specifically, after stimulus offset, participants first made a movement from the home position to one of two lateral targets based on their perceptual decision, and then made a lead-in movement to the via-point before moving to the end target. The force field (or channel) was applied between the via-point and the end target. The movements between the home position to lateral targets and from the lateral targets to the via-point were always made in a null field. Thus, by replicating the choice-before-movement structure of Experiment 2a with a distinct motor structure, this experiment allowed us to test whether the effects observed in Experiment 2a generalized beyond the specific geometry of the follow-through paradigm. It also provided a stronger test of advance commitment than button choice, as unlike button choice, the lead-in paradigm removes the possibility of revising the chosen action after the force field is experienced.

### Miss trials and incomplete dataset across experiments

Trials were missed if participants did not execute movements fast enough or they did not respect the trial structure of the experiments. In these cases, the trial was immediately aborted and a feedback messages was displayed depending on the type of miss trial. A “move sooner” message was displayed if movement was not initiated within 1 s of the auditory cue in Experiments 1a–b. A “decide sooner” message appeared if participants failed to press a button of the response box in Experiment 1c & 2a or failed to select a target in Experiment 2b within 0.5 s of stimulus disappearance. We also monitored movement timing: in Experiments 1a–c & 2a, a “too slow” message was displayed if movement exceeded 0.5 s from home position to via-point or 1 s from via-point to the lateral end targets, whereas in Experiment 2b it was displayed if movement exceeded 1 s from the lateral targets to the via-point or 0.5 s from via-point to the end target. In Experiment 2b, a “make a choice first” message was displayed if participants moved directly to the via-point without making a prior choice. Finally, in all experiments, a “you hit the wall” message was displayed if the cursor contacted the wall during movement through the passage. If a miss trial occurred, that trial was immediately repeated with the same coherence to preserve the planned block structure. To exclude the possibility that these repeated trials might have biased our results, we performed all data analyses on the testing phase by omitting these trials (3.5% of trials).

Two participants (one each from Exps. 1c and 2b) completed only 91% and 82% of the experiment, respectively, due to technical issues. However, we retained these participants in the analysis because they still contributed sufficient testing-phase data; the missing data corresponded to 3 of 12 and 6 of 12 channel trials per coherence, respectively.

### Data analysis

We used channel trials to compute adaptation as the proportion of the force field that was compensated for by regressing the actual forces, *f* (*t*), generated by participants in the channel against the ideal forces, *f* ^∗^(*t*), that would have fully compensated for the forces on a force-field trial (Eq. 1) [20, 21]. That is, we estimated *a*, the regression coefficient reflecting our measure of adaptation, from *f* (*t*) = *a f* ^∗^(*t*), where *t* is the discrete time step in the trial. (Note that the offset of the regression was constrained to zero.) For this analysis, we only used the portion of the movement during which hand velocity exceeded 1 cm s^−1^ within the channel.

#### Model-free analyses

During the testing phase, we quantified perceptual sensitivity to motion coherence using a model-free psychometric analysis. For each participant, we fit a probit regression to choice data from channel trials, with binary choice outcome (rightward vs. leftward) as the dependent variable and signed motion coherence *C* as the predictor. Perceptual sensitivity was quantified as the slope of the fitted psychometric function at its point of subjective equality.

If contextual uncertainty leads to the mixing of memories then the magnitude of adaptation expressed should be greatest for *C* = ±1 and smallest for *C* = ±0.1. To test this hypothesis, we fit a linear regression model to adaptation *a* during the testing phase. Critically, to avoid the confound of seemingly graded adaptation caused purely by a graded fraction of trials in which the participant judged the stimulus to be left or right, we fitted left and right choice trials separately. Specifically, we fitted separate intercepts (*α*^±^ at *C* 1) to right and left choice trials but, to minimize the number of fitted parameters, we assumed the effect of *C* was the same for both choices (i.e. same slope, *β*):

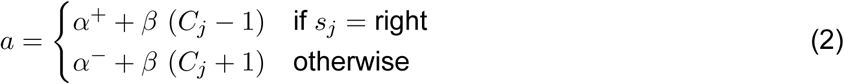

As our main interest was the dependence of adaptation on coherence, in order to increase the sensitivity of our analyses in the face of substantial variability in the overall amount of adaptation at *C* = ±1 coherence across participants (Suppl. Fig. 2), we computed normalized slopes

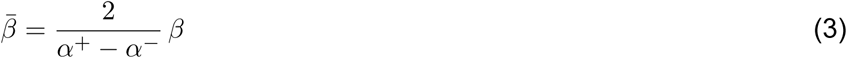

To plot data across participants, we normalized both data and the regression lines by shifting them by 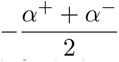 and then scaling them by 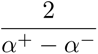 so that the intercepts of the regression lines for right and left choice trials at *C* ± 1 were normalized to be at ±1, respectively. All statistics reported for these plots (see below) refer to these normalized data and regression lines. (Note that the regression slopes shown on these normalized plots are identical to the normalized slopes of Eq. 3.)

We quantified evidence for a non-zero mean normalized slope across subjects using both a one-sample *t* test and a JZS Bayes factor. For each group, we computed the one-sample *t* statistic against zero and converted this to BF_10_ under a Jeffreys–Zellner–Siow prior with scale *r* = 2/2. The Bayes factor was obtained by numerically evaluating the marginal likelihood under the alternative hypothesis and dividing it by the marginal likelihood under the null. Values BF_10_ *>* 1 indicate evidence for a non-zero slope, whereas BF_10_ *<* 1 indicate evidence in favor of the null.

We compared slopes between experiments using two-sample Welch’s t-tests and the degrees of freedom were estimated using the Welch-Satterthwaite approximation.

#### Model-based analyses

We also fit two Bayesian ideal observer models to each experiment (Fig. 3). Both of these models infer context based on the RDK stimulus. In the first model, in line with the predictions of the contextual inference (COIN) model of motor adaptation [7], both memories are expressed and additively combined, each weighted by the probability with which it is inferred to be relevant in the current trial – that is memory combination (*MC*). In the second model, only the most probably relevant memory is expressed – that is memory selection (*MS*).

According to the model, right (*D* = right) and left contexts (*D* = left) are sampled with prior probabilities

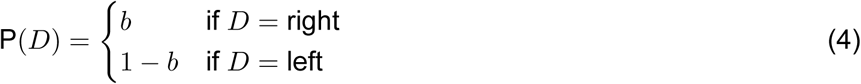

(where *b* stands for “bias”). As a partial mismatch to the true generative model of the experiment (see also below), the model assumes that the distribution of coherences is continuous (and unbounded):

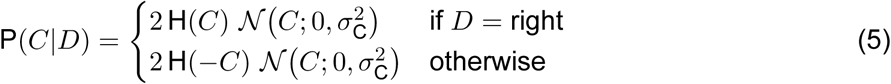

where H( ) is the Heaviside function, so that the resulting distributions are truncated Gaussians. Finally, the model assumes that the noisy sensory observation *x* is drawn from a normal distribution with mean *C* (coherence), and standard deviation *σ*_obs_:

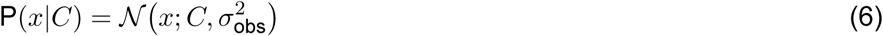

(This corresponds to the decision variable in an unbounded drift-diffusion model with fixed trial duration.) Thus, by marginalizing over *C*, we obtain the probability of observing *x* under *D* = right (right context) or *D* = left (left context), respectively, as:

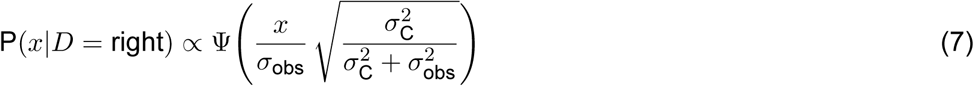

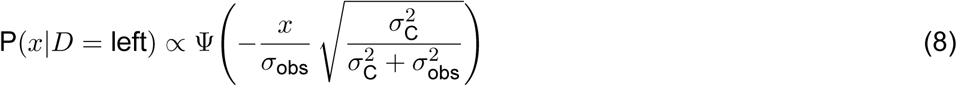

where Ψ(•) is the standard normal c.d.f., and we suppressed the constants of proportionality that only depend on *x* but not on *D* (and thus do not affect the inferences of the observer). Taking the limit of *σ*^2^ → ∞, we obtain the final forms of the likelihood:

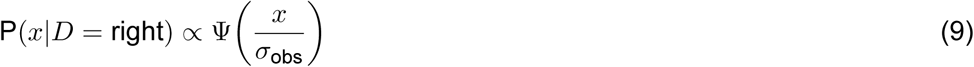

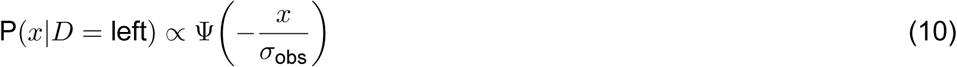

With these, the posterior beliefs over contexts are:

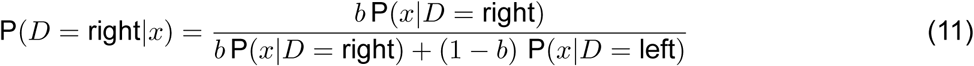

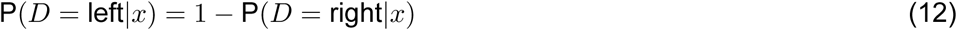

and we assumed that the same posterior over *D* drives both motor adaptation (using either memory combination or memory selection in the expression of motor memories, see below) and perceptual decisions.

We note that we also considered an alternative variant for Eq. 5, in which the observer’s internal model was fully matched to the true generative model of the experiment, such that coherences are sampled according to their experimentally defined distribution (and all other parts of the model, Eqs. (4) and (6), are unchanged):

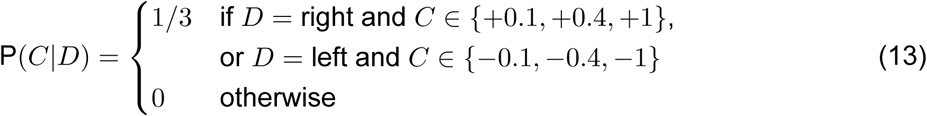

in which case the final forms of the likelihood (the equivalents of Eqs. (9) and (10)) are trivially:

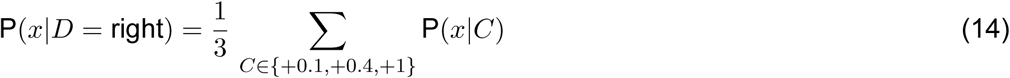

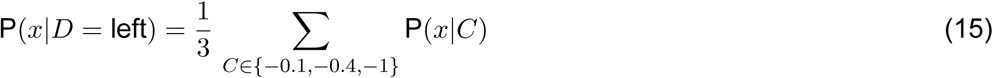

and the posterior is still computed as in Eqs. (11) and (12). As quantitative model comparison (ΔBIC = 15.81 across all participants, see below) revealed that this model provided a slightly poorer fit to experimental data than the mismatched model described above (results not shown), we did not consider it any further.

### Motor adaptation: memory combination (MC)

According to memory combination, the level of motor adaptation, *a*, is determined as a mixture of motor memories, such that each motor memory is weighted by the posterior probability with which its corresponding context is believed to be relevant in a given trial:

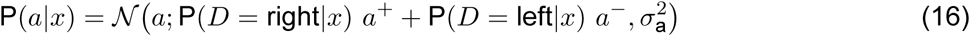

where *a*^−^ and *a*^+^ are the learned memories for contexts *D* = left and *D* = right, respectively, and *σ*^2^ is the variance of motor noise. (Combining memories by a simple linear mixture, as here, is normative when the loss function can be assumed to be quadratic as a function of motor adaptation [7].)

### Motor adaptation: memory selection (MS)

According to memory selection, only the most probably relevant memory is expressed on every trial:

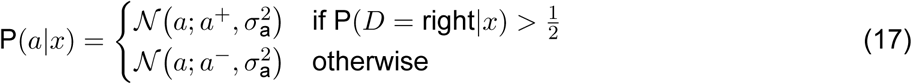

To simplify notation later, we unify Eqs. (16) and (17) using a softmax approach:

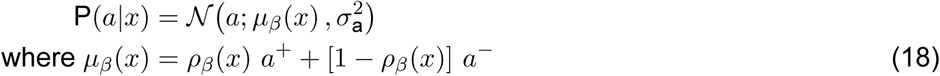

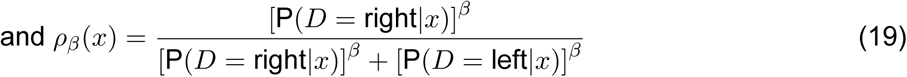

with *β* = 1 for memory combination and *β* → ∞ for memory selection.

### Perceptual decision

Binary perceptual decisions were modeled as reflecting the most probable context, without any decision noise:

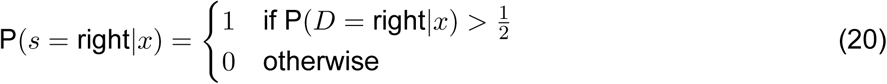

where *s* is the decision.

### Model fitting and comparison

For each participant, we fit each model (our model of perceptual decisions combined with either the combination or the selection model of motor memory expression, see above) to the behavioral observations produced by the participant, = *a_j_, s_j_* , including their adaptations, *a_j_*, and choices, *s_j_*, where *j* indexes the channel trials (both correct and incorrect) of the testing phase. To fit a model, we minimized its negative log-likelihood using fminsearchbnd (choosing the best fit from 32 random initial starts):

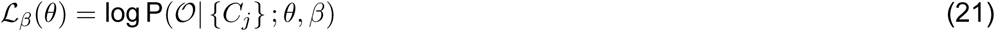

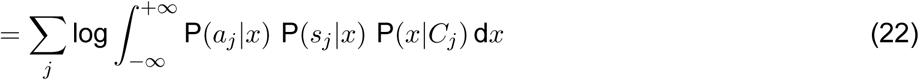

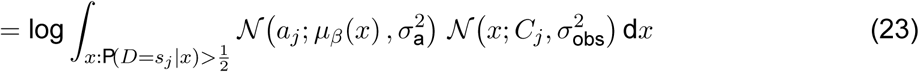

where *β* = 1 for memory combination and *β* for memory selection (as in Eq. 19), *θ* is the set of free parameters *a*^−^*, a*^+^*, σ*_obs_*, b, σ*_a_ , and note that P(*D* = *s_j_ x*) and *ρ_β_*(*x*) also depend on *σ*_obs_ and *b* (we just suppressed these dependencies for notational convenience).

Model comparison was performed using BIC, computed separately for each participant for each model. Since the models have the same number of parameters:

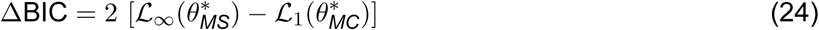

where 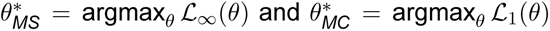 are the best fit parameters of the *MS* and *MC* models, respectively.

A ΔBIC above +6 or below 6 is considered to provide strong evidence in favor of the *MS* or *MC* model, respectively [48]. To perform model comparison at the group level, we calculated the group level ΔBIC as the sum of ΔBIC over individuals [49].

### Parameter and model recovery

We used the parameters from the fits of the *MC* and *MS* models to the data of each participant to generate five synthetic datasets per model class for each participant. For parameter recovery (Suppl. Fig. 3a–j; all *R*^2^ *>* 0.82), we used the parameters that were used to generate the synthetic data (true parameters) with the fit parameters to the synthetic datasets (recovered parameters) for the same model class. For model recovery (Suppl. Fig. 3k), we fit both model classes to each synthetic dataset as we did with the real data (see above). Thereafter, we examined the proportion of times that ΔBIC favored the true model class that generated the data (91% for *MC* and 81% for *MS*).

### Plotting model predictions

For plotting predicted psychometric curves, we computed

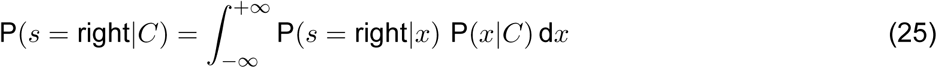

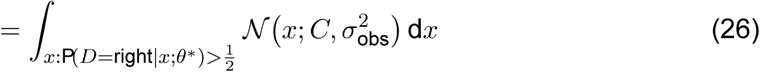

using the the best-fit parameters (maximizing Eq. 23).

For plotting predicted adaptations, we computed

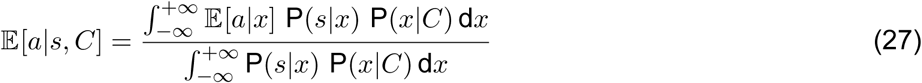

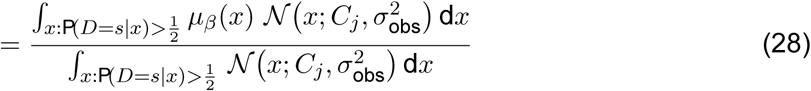

where E[·] denotes expected value.

For plotting average results across participants, we normalized both data and the model’s predictions analogously to the normalization procedure used in the model-free analyses (see above), but using *a*^+^ and *a*^−^ instead of *α*^+^ and *α*^−^, respectively.

## Data Availability

Data will be made available on Mendeley.

## Code Availability

Code will be made available on Mendeley.

## Acknowledgments

This work was supported by the grant SFI-AN-NCSCN-00007276 from the Simons Foundation International as part of the SCENE collaboration (to D.M.W and M.L.) and a Wellcome Trust (Investigator Award in Science 212262/Z/18/Z to M.L.).

## Author Contributions

All authors conceived the experiments and wrote the manuscript. A.N. conducted the experiments and analyzed the data.

## Competing Interests

The authors declare no competing interests.

---

**Supplementary Fig. 1.**
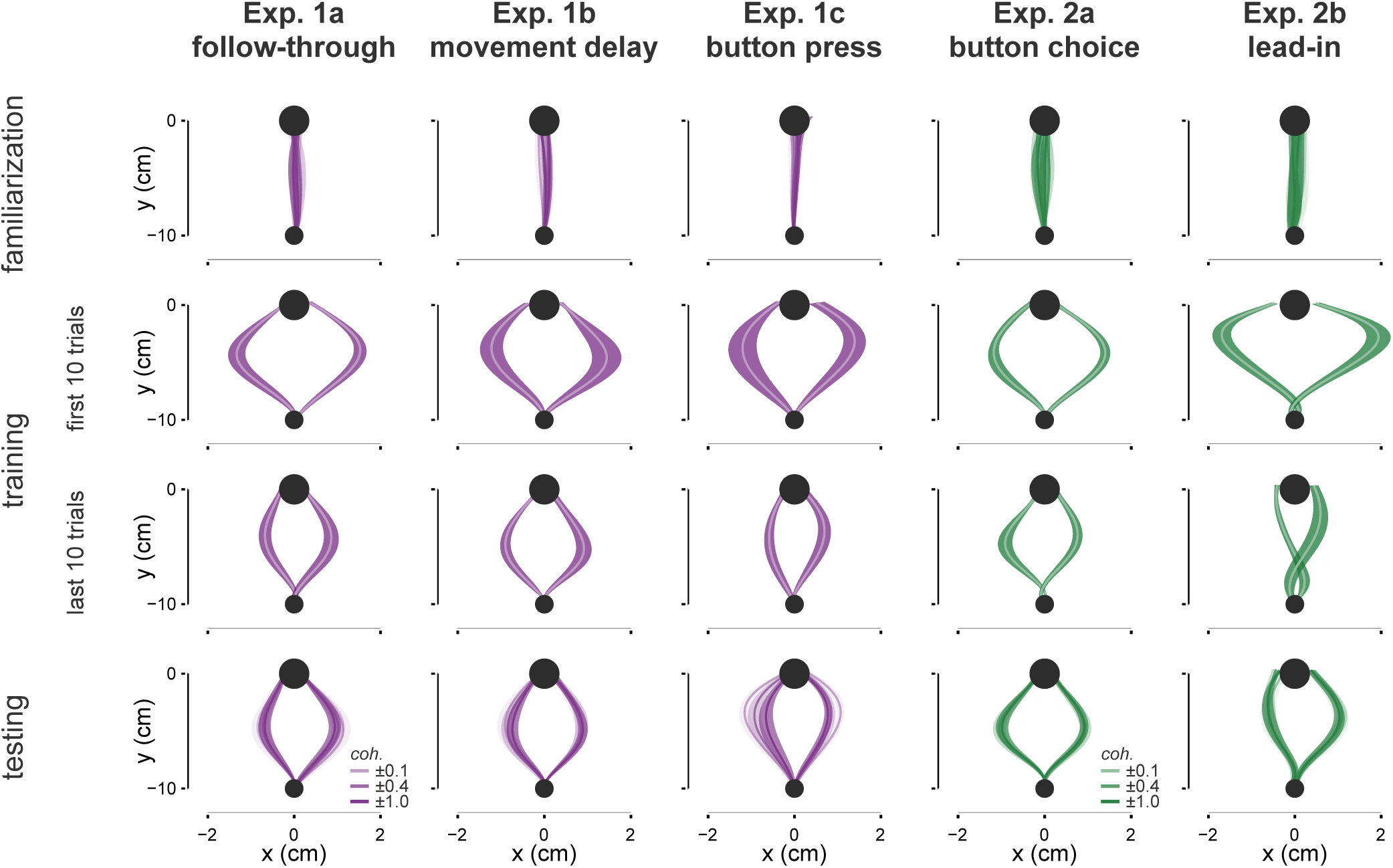
Progression of movement paths. Movement paths during non-channel trials in all experiments (columns). Average path for all trials of the familiarization trials (first row), the first 10 training trials (second row), last 10 training trials (third row) and all trials of testing phase partitioned by coherence (fourth row). Paths show mean *±* s.e.m. across participants.

**Supplementary Fig. 2.**
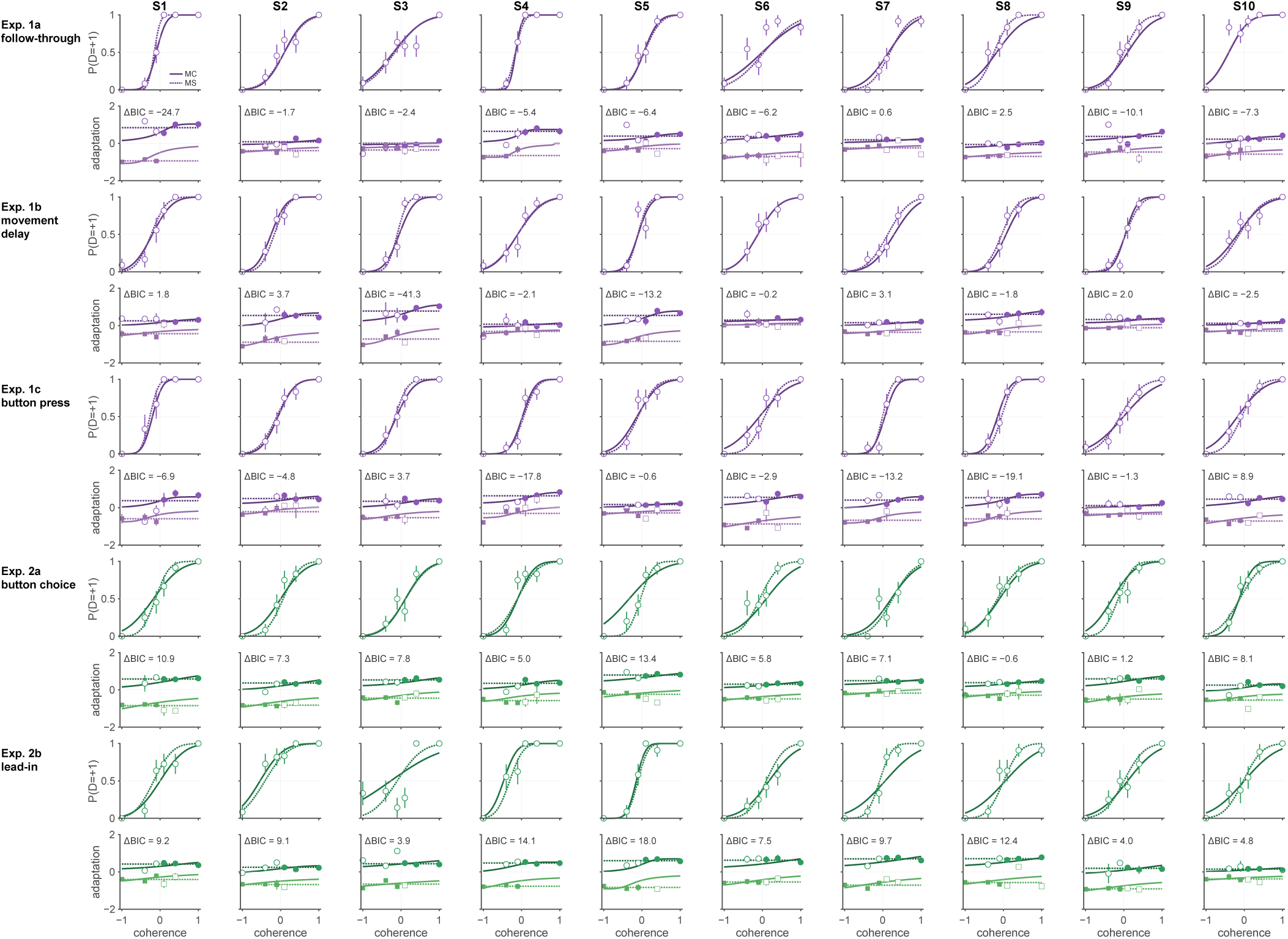
Bayesian observer model fits to adaptation to individuals. Probability of rightward choice (first row) and fits to adaptation (second row) with *MC* and *MS* models (as in Fig. 4a,b) for each experiment (rows) and participant (columns). ΔBIC for individuals with negative in favor of *MC* are shown in each plot.

**Supplementary Fig. 3.**
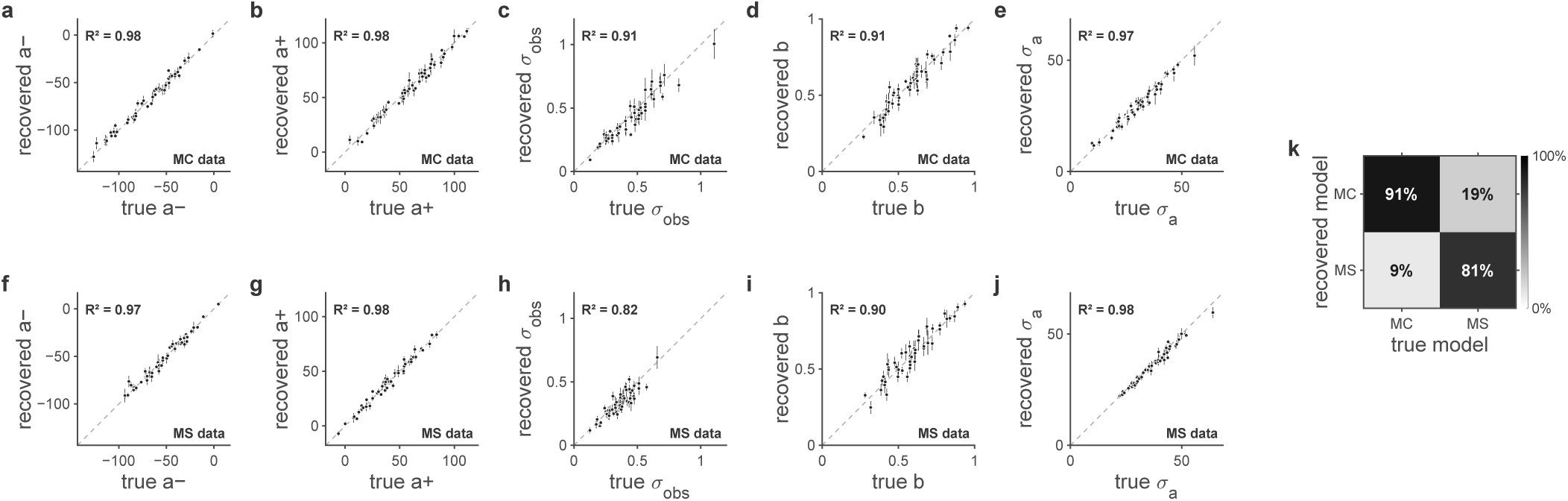
Parameter and model recovery. Parameter recovery of the five parameters for all experiments for **a–e.** *MC* and **f–j.** *MS* synthetic data (mean *±* s.e.m. across the 5 datasets generated for each participant). **k.** Model recovery for all experiments.

## Notes

### Competing Interest Statement

The authors have declared no competing interest.

